# Modular counterplays shape condensation and signaling of chimeric antigen receptor

**DOI:** 10.64898/2026.07.27.741008

**Authors:** Yiran Jiang, Zhengxu Ren, Jijun Luo, Haochen Yang, Xiwei Liu, Hui Chen, Chenqi Xu, Jizhong Lou

## Abstract

Chimeric antigen receptor-engineered T (CAR T) cell therapy has achieved remarkable clinical efficacy against hematological malignancies but exhibits limited therapeutic outcomes in solid tumors, where immune checkpoint signaling is highly active. Here, we demonstrate that the second-generation CD28-CD3ζ (28Z) CAR undergoes liquid-liquid phase separation with the Src-family kinase LCK to assemble a signaling condensate crucial for CAR activation. PD-1, but not LAG3 or CTLA4, disrupts these CAR-LCK condensates through competitive binding to CAR rather than through its canonical phosphotase-dependent inhibitory pathway. Incorporation of a CD3ε cytoplasmic module into the CAR introduces additional LCK interactions that reinforce CAR-LCK condensation, thus limited PD-1 incorporation and stabilize signalosome assembly. Consequently, the CD3ε-engineered CAR (E-CAR) resists PD-1-mediated condensates disassembly, preserves immunological synapse organization, and sustains proximal signaling following PD-L1 engagement. E-CAR T cells are therefore resistant to PD-1-mediated functional suppression and maintain potent antitumor activity against PD-L1-positive solid tumors. Together, these findings identify disruption of CAR signaling condensates as a previously unrecognized mechanism of PD-1-mediated inhibition and establish CD3ε modular signalosome engineering as a rational engineering strategy to overcome immune checkpoint suppression.

## Introduction

CAR-T cell therapy has emerged as a transformative targeted immunotherapeutic modality for hematologic malignancies ^1–3^. Nevertheless, accumulating clinical and preclinical evidences has documented its poor response durability and limited antitumor persistence against solid tumors^4^. A primary driver of CAR-T therapeutic failure and tumor relapse is the immunosuppressive tumor microenvironment (TME), with multiple inhibitory molecules reshape TME homeostasis, induce CAR-T cell exhaustion^5,6^, and directly or indirectly perturb CAR proximal signaling and co-stimulation cascades^7,8^, thereby drastically abrogating CAR-T effector functions. These translational bottlenecks necessitate mechanistic dissection of CAR-T signaling regulation and rational design of solid-tumor-adapted CAR constructs.

Immune checkpoints are a series of inhibitory receptors expressed on the surface of immune cells. Under the stimulation of ligands, they activate downstream inhibitory signaling pathways, thereby inhibiting the function of immune cells. Immune checkpoint molecules play a key role in the immune escape of tumors^9^. The PD-1/PD-L1 immune checkpoint axis represents a dominant inhibitory pathway^2,10–14^, that restrains endogenous ^15^ and CAR-T antitumor immunity^16–19^, enabling tumor immune evasion. Canonically, PD-1 ligation recruits phosphatases to dismantle T cell signalosomes, which constitutes a core barrier to effective antitumor immune responses. Clinically, the anti-tumor function of immune cells can be restored by blocking the binding of immune checkpoints to ligands, thereby achieving the purpose of treating tumors^20–22^. Cancer immunotherapy mainly based on PD-1/PD-L1 blocking antibodies has achieved great success in clinical practice^5,21,23^. While PD-1 blockade therapies targeting PD-1 phosphorylation and subsequent SHP2 recruitment yield durable remission in subsets of patients, heterogeneous treatment responses remain poorly mechanistically explained^24,25^. Additionally, LAG3 has emerged as another important inhibitory checkpoint for cancer immunotherapy. Recent studies further revealed that LAG3 suppresses T cell activation by disrupting CD3ε-LCK condensates or promoting TCR degradation^26,27^. Whether these inhibitory checkpoint pathways similarly regulate CAR signalosome assembly and function, however, remains largely unknown.

Biomolecular condensation governs diverse cellular physiological processes^28–34^. Recent studies have demonstrated that CAR molecules undergo phase separation upon antigen engagement, promoting CAR clustering, immunological synapse formation, antigen sensitivity, and T cell persistence^35,36^. However, these studies have primarily focused on CAR self-condensation, whereas the molecular mechanisms governing assembly of the CAR signaling condensate remain largely unknown. Notably, CD28 has recently been shown to directly condensate with the Src-family kinase LCK, thereby facilitating signal initiation.^25,37^ How CAR signaling condensates recruit and organize LCK during activation therefore remains an unresolved question. In parallel, CAR-T cells rapidly upregulate PD-1 following activation^38^, while PD-L1 is abundantly expressed by both malignant cells and stromal components within the tumor microenvironment. and widespread PD-L1 expression on malignant parenchymal and stromal cells within the TME^39^, Consequently, CAR signaling condensates are continuously exposed to inhibitory checkpoint signaling *in vivo*. Whether PD-1, as well as other inhibitory regulators such as LAG3, directly remodels CAR signaling condensates, and how engineered CARs overcome such inhibition, remain unknown.

Here, we established an *in vitro* phase-separation system to reconstitute CAR signalosome assembly and uncovered the modular rules governing the CAR-LCK condensate formation. We show that PD-1 directly disrupts CAR signalosome assembly independently of canonical phosphatase signaling, whereas incorporation of the CD3ε intracellular domain introduces an additional LCK-binding module that stabilizes condensate assembly and counteracts PD-1 incorporation. Consequently, E-CAR preserves immunological synapse organization and antitumor function despite PD-1/PD-L1 engagement. Together, our findings establish a modular competition model for signalosome assembly, in which the composition of signaling modules determines interaction partner selection and functional output, providing a rational framework for engineering immune checkpoint-resistant CAR-T cells.

## Result

### Molecular condensation between 28Z CAR and LCK is selectively inhibited by PD-1 cytoplasmic tail

To biochemically reconstitute CAR activation, we purified the cytoplasmic tails of 28Z and BBZ CARs and phosphorylated them by LCK kinase domain *in vitro*. Mass spectrometry confirmed extensive phosphorylation of all predicted tyrosine residues within the 28Z intracellular domain **(Supplementary Table 1)**. We next examined condensate formation of phosphorylated CARs cytoplasmic tails together with LCK_UD-SH3-SH2_ on supported lipid bilayers (SLBs), in short LCK-CAR condensation, which recapitulate the membrane environment during CAR activation. Strikingly, phosphorylated 28Z (p28Z), but not phosphorylated BBZ (pBBZ), formed condensates with LCK **(Fig. 1A)**. Compared to the unphosphorylated CAR, its phosphorylation further enhanced condensate assembly, resulting in larger and denser condensates with increased LCK enrichment **(Fig. 1B)**. Notably, pre-formed p28Z condensates efficiently recruited unphosphorylated 28Z molecules **(Fig. 1C)**, suggesting that phosphorylation serves as a nucleation event to promote progressive signalosome assembly through phase separation.

**Figure 1.**
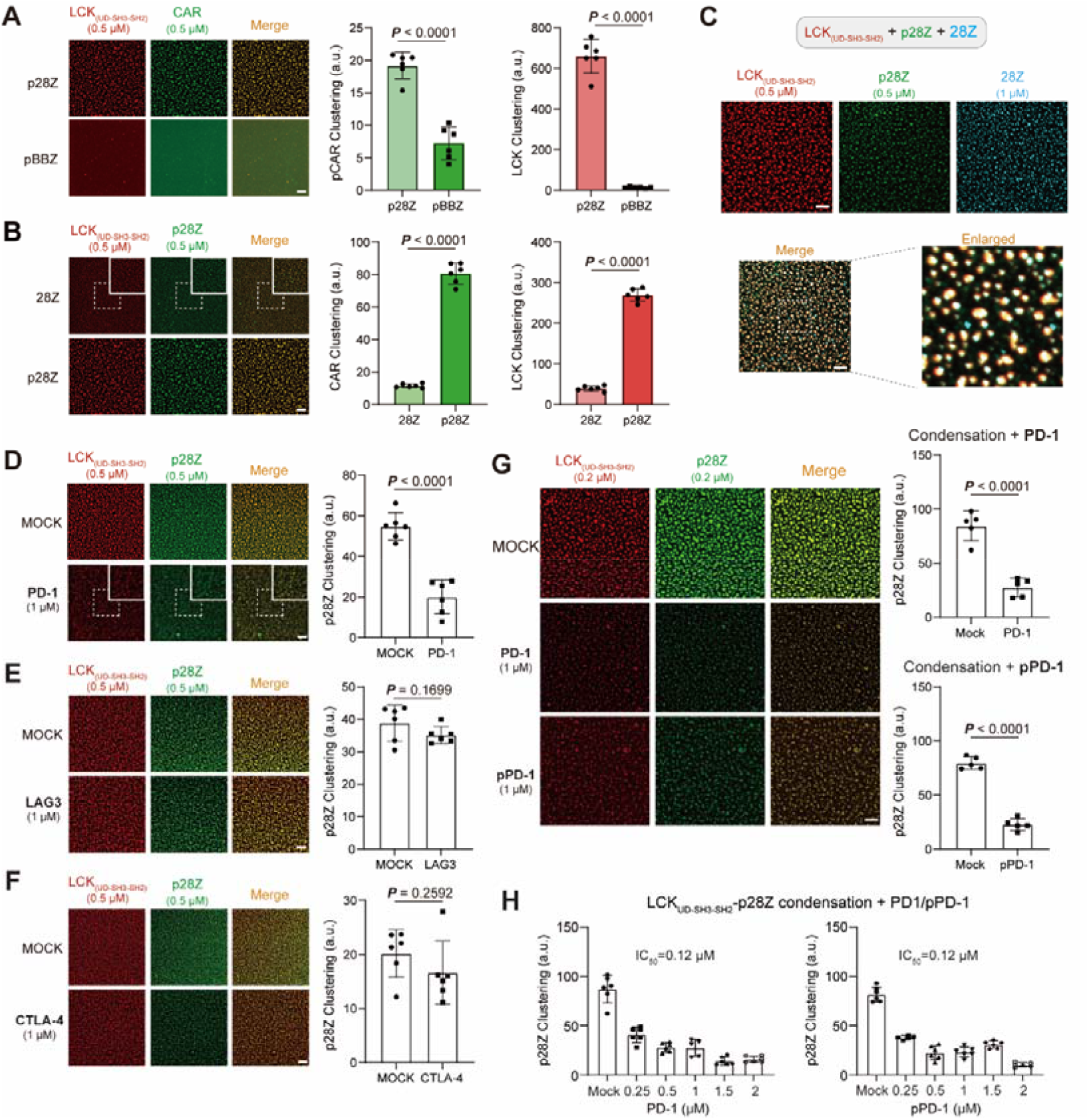
PD-1 but not LAG3 or CTLA4 inhibits LCK-p28Z CAR phase separation. **(A-B)** Membrane phase separation between phosphorylated CAR and LCK. The phosphorylated cytoplasmic tails of 28Z or BBZ CAR (p28Z or pBBZ) were generated by incubation with the LCK kinase domain. Fluorescently labeled LCK_UD-SH3-SH2_ at the indicated concentrations was immobilized on supported lipid bilayers (SLBs) via a His tag, followed by the addition of phosphorylated (A) or unphosphorylated (B) CAR cytoplasmic tails at the indicated concentrations. Samples were incubated at room temperature for 60 min before imaging. Condensation was quantified as described in the Methods (n = 6). Scale bar, 5 μm. **(C)** Effect of unphosphorylated 28Z on preassembled LCK_UD-SH3-SH2_-p28Z (in short LCK-p28Z in text) condensates. LCK_UD-SH3-SH2-p28Z_ condensates were first assembled as described in (A-B), followed by the addition of unphosphorylated 28Z cytoplasmic tail and incubation at room temperature for 30 min before imaging. Scale bar, 5 μm. **(D-F)** Effects of PD-1 (D), LAG3 (E), and CTLA-4 (F) cytoplasmic tails on LCK_UD-SH3-SH2_-p28Z condensates. LCK_UD-SH3-SH2_-p28Z condensates were preassembled as described in (A-B), followed by the addition of the indicated checkpoint receptor cytoplasmic tails at the indicated concentrations and incubation at room temperature for 30 min before imaging (n = 6). Scale bar, 5 μm. **(G-H)** Effects of PD-1 and phosphorylated PD-1 (pPD-1) on LCK UD-SH3-SH2-p28Z condensates. Experiments were performed as described in (D-F). Representative images are shown in (G) (n = 5). PD-1 or pPD-1 was added at the indicated concentrations (H), and IC values of (p)PD-1 were determined by fitting the data with a four-parameter logistic regression model in GraphPad Prism (n = 6). Scale bar, 5 μm. Data are presented as mean ± standard deviation (SD). Statistical analyses were performed using two-tailed unpaired t-tests.

As immune checkpoints suppress CAR-T cell function, we next examined whether their cytoplasmic tails directly regulate CAR signalosome assembly. Among the cytoplasmic tails tested, only PD-1 markedly disrupted LCK-p28Z condensates, whereas LAG3 and CTLA-4 had minimal effects **(Fig. 1D-F)**, indicating a specific role for PD-1 in regulating 28Z signalosome assembly. PD-1-mediated condensate disruption occurred in a dose-dependent manner, and phosphorylated PD-1 (pPD-1) exhibited a comparable inhibitory potency to its unphosphorylated counterpart **(Fig. 1G)**, with similar IC values, about 0.12 μM **(Fig. 1H, Extended Data Fig. 1A and 1B)**. Together, these findings demonstrate that phosphorylation drives 28Z signalosome assembly, which is selectively disrupted by the PD-1 intracellular domain.

### PD-1 disrupts LCK-28Z CAR condensates through binding competition

To investigate how PD-1 disrupts LCK-p28Z condensates, we compared the ability of PD-1 and LCK to assemble condensates with p28Z over a range of protein concentrations on SLBs. At a fixed concentration of 0.1 μM PD-1 or pPD-1, 0.5 μM p28Z was sufficient to form robust condensates (**Fig. 2A-2C**). In contrast, under the same conditions, 0.1 μM LCK did not phase separate with p28Z (**Fig. 2C-2E**), whereas at least 0.2 μM LCK was required to support robust condensate formation with 0.5 μM p28Z (**Fig. 2C and 2E**). These results indicated that PD-1 associated with p28Z more efficiently than LCK, enabling PD-1 to competitively displace LCK from p28Z and thereby disrupt LCK-p28Z condensates.

**Figure 2.**
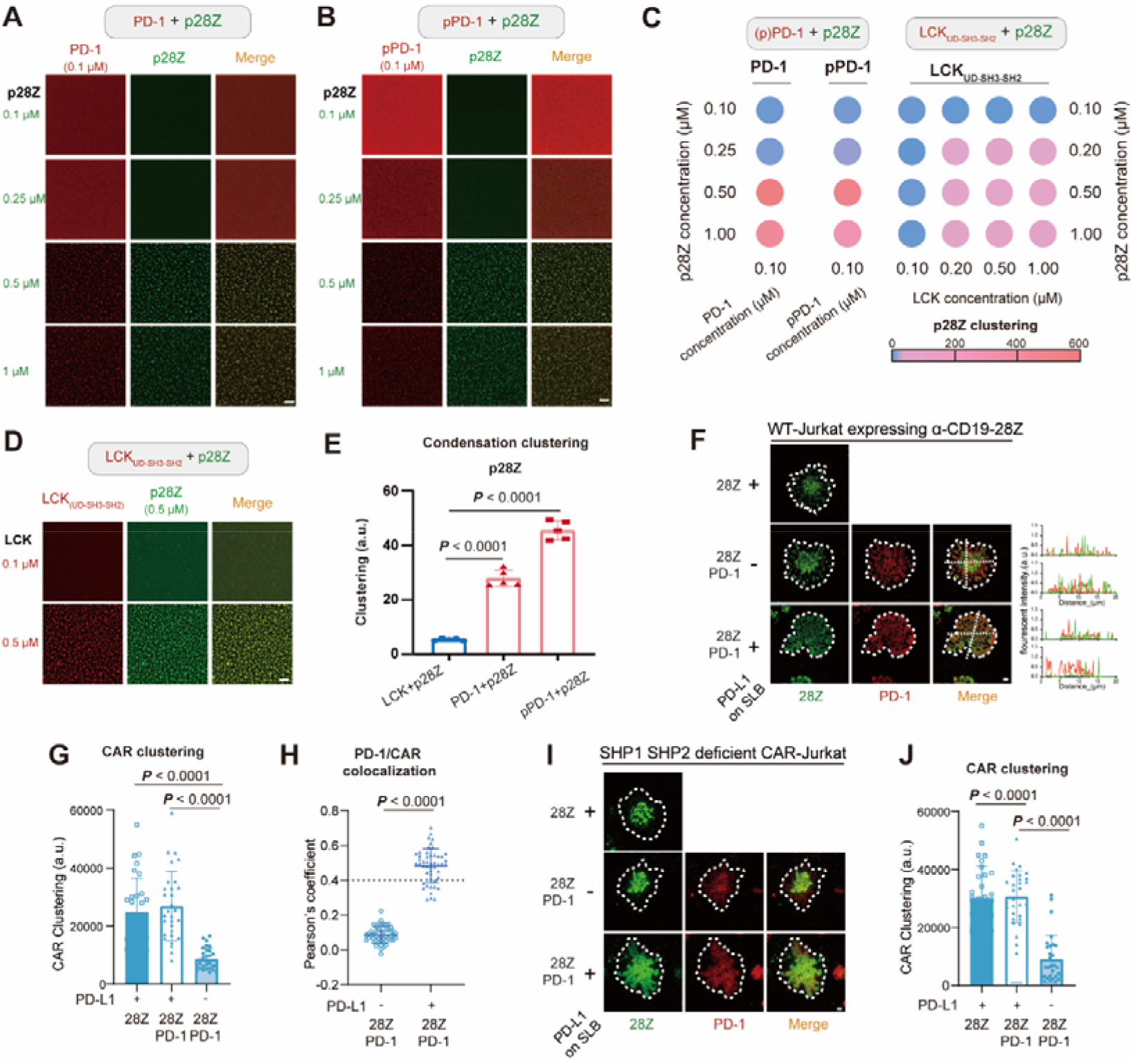
PD-1 disrupts CAR-LCK signaling condensates and impairs immunological synapse assembly independently of SHP1/2 phosphatases. **(A-B)** On-membrane condensation between p28Z and PD-1 (A) or phosphorylated PD-1 (pPD-1) (B). Experimental procedures are described in the Methods. PD-1 or pPD-1 was held constant at 0.1 μM, whereas p28Z was added at the indicated concentrations. Scale bar, 5 μm. **(C)** Heatmap showing p28Z clustering of PD-1-p28Z, pPD-1-p28Z, and LCK_UD-SH3-SH2_-p28Z condensates assemblies across the indicated protein concentrations. PD-1 or pPD-1 was maintained at 0.1 μM, whereas LCK concentration was varied from 0.1 to 1 μM. Each protein was incubated with p28Z at the indicated concentrations, and p28Z clustering in condensates was quantified using Fiji (n = 5). **(D)** Representative images of LCK _UD-SH3-SH2_-p28Z condensates formed at the indicated protein concentrations. Scale bar, 5 μm. **(E)** Quantification of condensate clustering for PD-1-p28Z, pPD-1-p28Z, and LCK_UD-SH3-SH2_-p28Z assemblies formed at the same protein concentrations (0.1 μM PD-1/pPD-1/LCK_UD-SH3-SH2_ +0.5 μM p28Z). Condensation was quantified from the PD-1, pPD-1, or LCK_UD-SH3-SH2_ fluorescence channel using Fiji (n = 5). **(F-H)** Immunological synapse imaging of anti-CD19 28Z CARs in wild-type Jurkat cells. Jurkat cells expressing EGFP-tagged anti-CD19 28Z, E28Z, or E_B6I_28Z CARs together with PD-1-mScarlet3 were stimulated on supported lipid bilayers (SLBs) coated with CD19 (10 nM) in the presence or absence of PD-L1 (10 nM) at 37 °C for 1 h, followed by TIRF-SIM imaging. (F) Representative images of immunological synapses. Scale bar, 1 μm. (G) CAR clustering within the immunological synapse, quantified as SD²/mean using Fiji. (H) Colocalization of PD-1 and CAR molecules within the immunological synapse in the presence or absence of PD-L1 on SLBs. Colocalization was quantified using Pearson’s correlation coefficient calculated with Fiji (n = 30). **(I-J)** Immunological synapse imaging of anti-CD19 28Z CARs in SHP1/SHP2 double-knockout Jurkat cells. Imaging was performed as described for panels (F-H). **(I)** Representative images of immunological synapses. Scale bar, 1 μm. **(J)** CAR clustering within the immunological synapse, quantified as SD²/mean using Fiji. Data are presented as mean ± standard deviation (SD). Statistical analyses were performed using two-tailed unpaired t-tests.

To determine whether PD-1 regulates CAR signalosome assembly under physiological conditions, we visualized immunological synapse formation in Jurkat cells expressing anti-CD19 28Z CAR on CD19-containing supported lipid bilayers (SLBs) by total internal reflection fluorescence structured illumination microscopy (TIRF-SIM). Compared with control cells, PD-1 overexpression alone did not appreciably affect the frequency of mature immunological synapse formation or CAR microcluster assembly. Instead, PD-1 was predominantly distributed around the central CAR cluster. Upon PD-L1 engagement, however, PD-1-expressing CAR Jurkat cells exhibited markedly impaired synapse maturation (**Fig. 2F** and CAR clustering (**Fig. 2G**), accompanied by dispersion of the central CAR structure and increased colocalization of PD-1 with CAR (**Fig. 2H**). To determine whether this effect required canonical PD-1 downstream signaling, we performed the same imaging analysis in SHP1/SHP2 double-knockout Jurkat cells. Strikingly, PD-L1 still disrupted immunological synapse organization and CAR clustering to a similar extent in the absence of both phosphatases (**Fig. 2I and 2J**), demonstrating that PD-1 suppresses CAR signalosome assembly independently of SHP1/2-mediated downstream signaling.

### Addition of CD3**ε** module renders 28Z CAR resistant to PD-1-mediated condensate disruption

In the native TCR-CD3 complex, CD3ε recruits LCK through electrostatic interactions, and we previously showed that incorporation of the CD3ε cytoplasmic domain into 28Z CAR (E28Z) markedly enhances CAR self-condensation through cation-π interactions^35^. Because E28Z undergoes rapid endocytosis that limits its surface expression, we further optimized the construct by replacing the region between the basic-rich sequence (BRS) and ITAM with six glycine residues, generating E_B6I_28Z (E28Z and E_B6I_28Z CAR collectively referred to as E-CAR), which exhibited improved stability while preserving its enhanced condensate-forming capacity.

We next asked whether CD3ε incorporation confers resistance to PD-1-mediated condensate disruption. E-CAR ICDs were also phosphorylated by the LCK kinase domain *in vitro*, and phosphorylation was confirmed by mass spectrometry **(Supplementary Table 1)**. LCK was fixed at 0.2 μM while PD-1 ICD was titrated into LCK-p28Z or LCK-pE-CAR condensates. Although PD-1 disrupted all condensates in a dose-dependent manner, E-CAR condensates were markedly more resistant than 28Z condensates, with substantially increased IC_50_ values (0.12 μM for p28Z, 0.46 μM for pE28Z, and 0.30 μM for pE_B6I_28Z). A similar trend was observed with phosphorylated PD-1, where E-CARs condensates remained significantly less susceptible to disruption (**Fig. 3A-C, Extended Data Fig. 1A-B and Extended Data Fig. 2A-B**).

**Figure 3.**
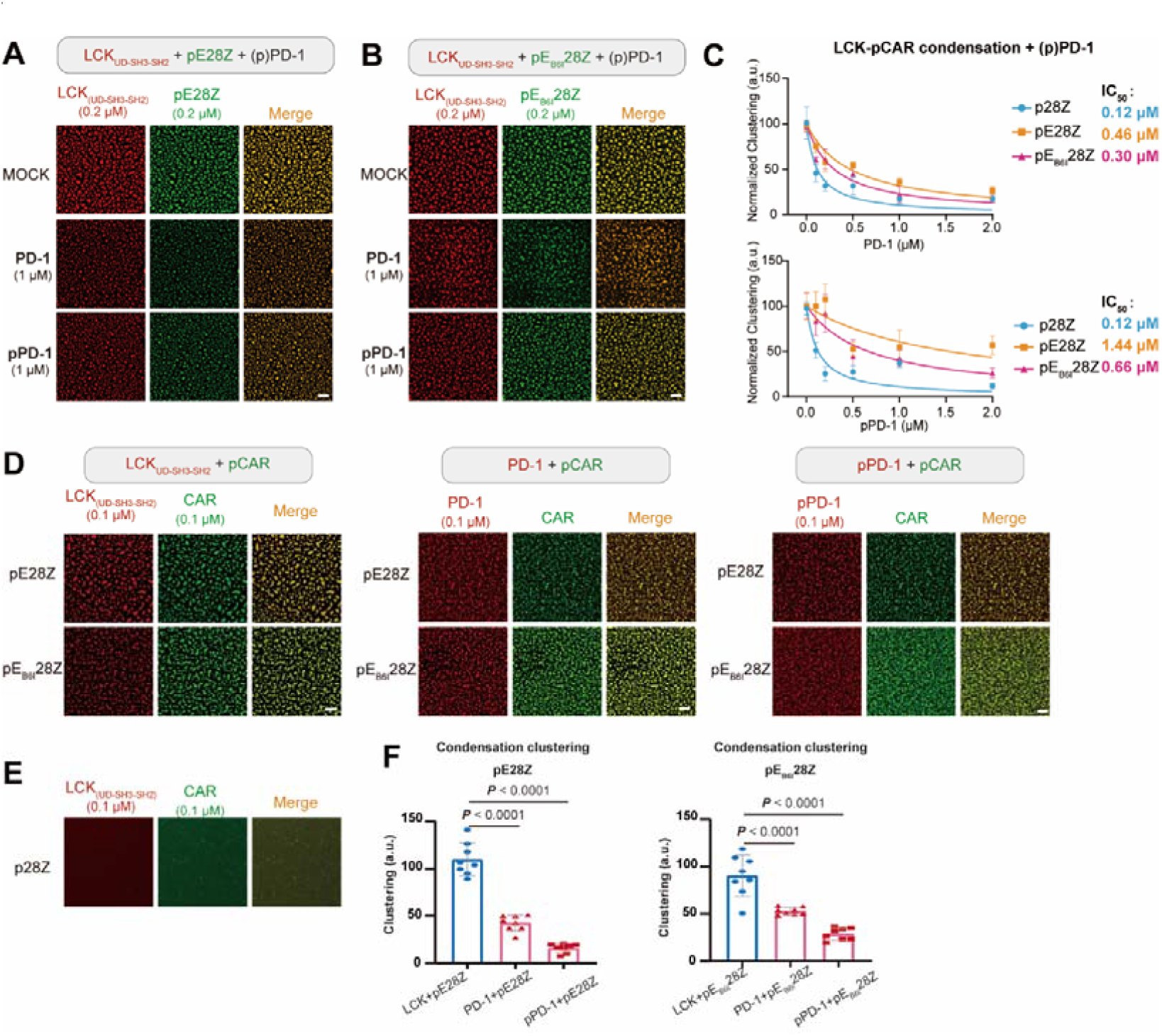
CD3ε provided additional LCK binding in E-CAR to form stronger condensation with LCK that resist to PD-1 competition. **(A-B)** Representative images of LCK_UD-SH3-SH2_-pE28Z (A) and LCK_UD-SH3-SH2_-pE_B6I_28Z (B) condensates following the addition of PD-1 or phosphorylated PD-1 (pPD-1) at the indicated concentrations. Experimental procedures were the same as those described in Figure 1D-G, in short LCK-pCAR condensates was pre-formed followed by adding PD-1 or pPD-1. Scale bar, 5 μm. (C) Quantification of LCK_UD-SH3-SH2_-pCAR condensate clustering in the presence of increasing concentrations of PD-1 or pPD-1. IC values were determined by fitting the data using a four-parameter logistic regression model in GraphPad Prism (n = 5). **(D-F)** Representative images (D and E) and quantification (F) of condensate clustering for PD-1-pE28Z, pPD-1-pE28Z, LCK_UD-SH3-SH2_-pE28Z, PD-1-pE_B6I_28Z, pPD-1-pE_B6I_28Z, and LCK_UD-SH3-SH2_-pE_B6I_28Z condensates formed at identical protein concentrations (0.1 μM PD-1, pPD-1, or LCK_UD-SH3-SH2_ together with 0.1 μM pCAR). Condensation was quantified from the PD-1, pPD-1, or LCK_UD-SH3-SH2_ fluorescence channel using Fiji (n = 8). Scale bar, 5 μm. Data are presented as mean ± standard deviation (SD). Statistical analyses were performed using two-tailed unpaired t-tests.

To understand the basis of this resistance, we compared the ability of LCK to assemble condensates with phosphorylated CARs under limiting LCK concentrations. At 0.1 μM LCK, both pE28Z and pE_B6I_28Z readily assembled robust condensates, whereas p28Z did not form condensates, indicating that CD3ε incorporation substantially enhanced LCK-driven condensate formation (**Fig. 3D-3F, Extended Data Fig. 3A**). In contrast, when PD-1 or phosphorylated PD-1 was added at the same concentration (0.1 μM), the resulting PD-1-containing condensates were markedly more weakly aggregated than those formed with LCK (**Fig. 3E and 3F**). These results suggested that incorporation of CD3ε shifted the competitive balance toward LCK, thereby stabilizing CAR signalosomes and rendering E-CAR condensates resistant to PD-1-mediated disassembly. As the CD3ε BRS has been reported to mediate selective LCK recruitment to the TCR-CD3 complex, we next examined whether the BRS similarly contributes to LCK recruitment in E-CAR. We generated a loss-of-function mutant (mE28Z), in which the positively charged lysine and arginine residues within the CD3ε BRS were substituted with alanine, while all tyrosine residues within the CD3ε ITAM were simultaneously mutated to phenylalanine to eliminate potential contributions from CD3ε phosphorylation (**Extended Data Fig. 3B**). Unlike wild-type E28Z, phosphorylated mE28Z failed to efficiently form condensates with LCK, even at 0.5 μM pmE28Z and 0.5 μM LCK, where p28Z and pE28Z phase separated with LCK. Notably, condensate formation was even weaker than that observed for LCK-p28Z (**Extended Data Fig. 3C and 3D**), suggesting that disruption of the CD3ε BRS abolished electrostatic LCK recruitment, while the additional nonfunctional CD3ε sequence may further reduce condensate formation through steric hindrance. Together, these results establish the CD3ε as a critical determinant of LCK recruitment and condensate assembly, thereby shifting the competitive interaction toward LCK over PD-1 and conferring resistance to PD-1-mediated signalosome disruption.

It has been reported that the LAG3 cytoplasmic tails and Csk disrupt LCK-CD3ε condensates during TCR activation by competing for binding to the CD3ε cytoplasmic tails^27,37,40,41^. We therefore asked whether incorporation of CD3ε would render E-CAR susceptible to LAG3- or Csk-mediated disruption. Unexpectedly, even at 1 μM, neither the LAG3 cytoplasmic tail nor Csk appreciably disrupted LCK-E-CAR condensates, with condensate clustering remaining comparable to the control (**Fig. 4A-4D**). One possible explanation is that the CD28 cytoplasmic tails provide additional LCK-binding interfaces, thereby strengthening LCK association with E-CAR and reducing its susceptibility to competition by both LAG3 and Csk. In addition, the CTLA-4 intracellular domain failed to affect LCK-E-CAR condensate assembly, even at 1 μM (**Fig. 4E and 4F**). Collectively, these findings demonstrate that E-CAR signalosomes remain resistant to multiple inhibitory regulators despite incorporation of the CD3ε cytoplasmic tails, highlighting the robustness of this engineering strategy in immunosuppressive environments.

**Figure 4.**
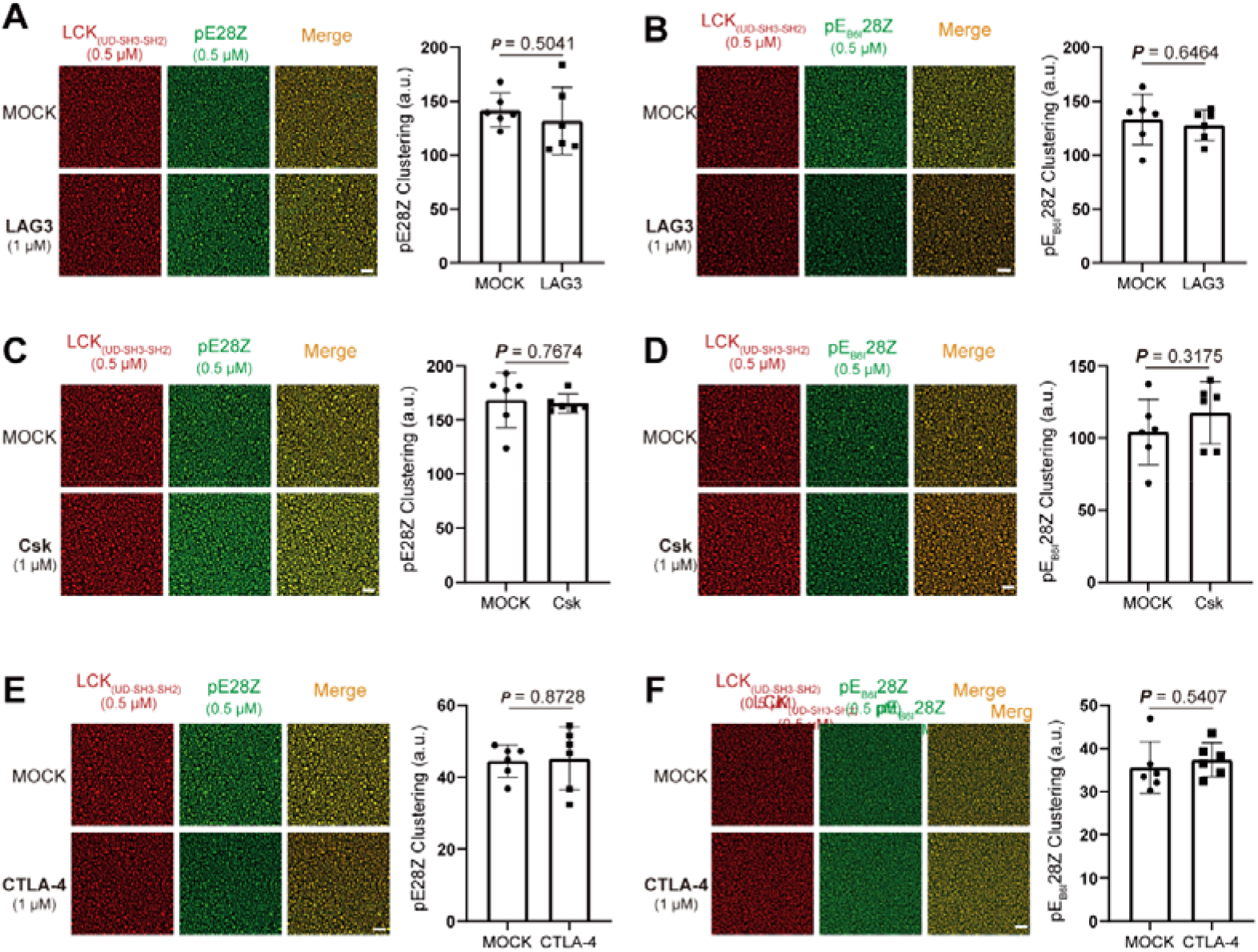
CD3ε engineering does not induce LAG3, Csk and CTLA-4 engagement in LCK-pE-CAR condensation. **(A-B)** Effect of LAG3 (1 μM) on preassembled LCK_UD-SH3-SH2_-pE28Z (A) and LCK_UD-SH3-SH2_-pE_B6I_28Z (B) condensates. Representative images and quantification of condensates clustering are shown. Experiments were performed as described in Figure 1D-G, in short LCK-pCAR condensate was pre-formed followed by adding LAG3. Scale bar, 5 μm. **(C-D)** Effect of Csk (1 μM) on preassembled LCK_UD-SH3-SH2_-pE28Z (A) and LCK_UD-SH3-SH2_-pE_B6I_28Z (B) condensates. Representative images and quantification of condensates clustering are shown. Experiments were performed as described in Figure 1D-G, in short LCK-pCAR condensate was pre-formed followed by adding Csk. Scale bar, 5 μm. **(E-F)** Effect of CTLA-4 (1 μM) on preassembled LCK_UD-SH3-SH2_-pE28Z (A) and LCK_UD-SH3-SH2_-pE_B6I_28Z (B) condensates. Representative images and quantification of condensates clustering are shown. Experiments were performed as described in Figure 1D-G, in short LCK-pCAR condensate was pre-formed followed by adding CTLA-4. Scale bar, 5 μm. Data are presented as mean ± standard deviation (SD). Statistical analyses were performed using two-tailed unpaired t-tests.

### PD-1 shows minimal inhibition of E-CAR synapse in T cells

Given that the cytoplasmic tails of E-CAR resisted PD-1-induced condensate disruption *in vitro*, we next asked whether E-CAR T cells could maintain immunological synapse organization following PD-1 ligation. Jurkat cells expressing CD19-targeting 28Z or E_B6I_28Z CARs were stimulated on CD19-functionalized supported lipid bilayers (SLBs) with or without PD-L1, followed by visualization by TIRF-SIM imaging. In 28Z CAR-T cells, PD-L1 ligation markedly reduced synapse size, disrupted central CAR clustering, and disorganized the characteristic bull’s-eye architecture of the immunological synapse (**Fig. 5A**). In contrast, E28Z and E_B6I_28Z CAR-T cells exhibited only modest changes upon PD-L1 ligation and largely preserved an intact bull’s-eye synapse architecture (**Fig. 5B-5D**). Consistent with this observation, E-CAR displayed reduced colocalization with PD-1 at the synapse (**Fig. 5E**), suggesting diminished incorporation of PD-1 into the signaling complex. Importantly, phosphorylation of LAT remained unchanged in E-CAR T cells following PD-L1 ligation (**Fig. 5F**), indicating that downstream signaling was largely preserved despite checkpoint engagement. Together, these findings demonstrate that E-CAR maintains immunological synapse integrity and downstream signaling following PD-1 ligation.

**Figure 5.**
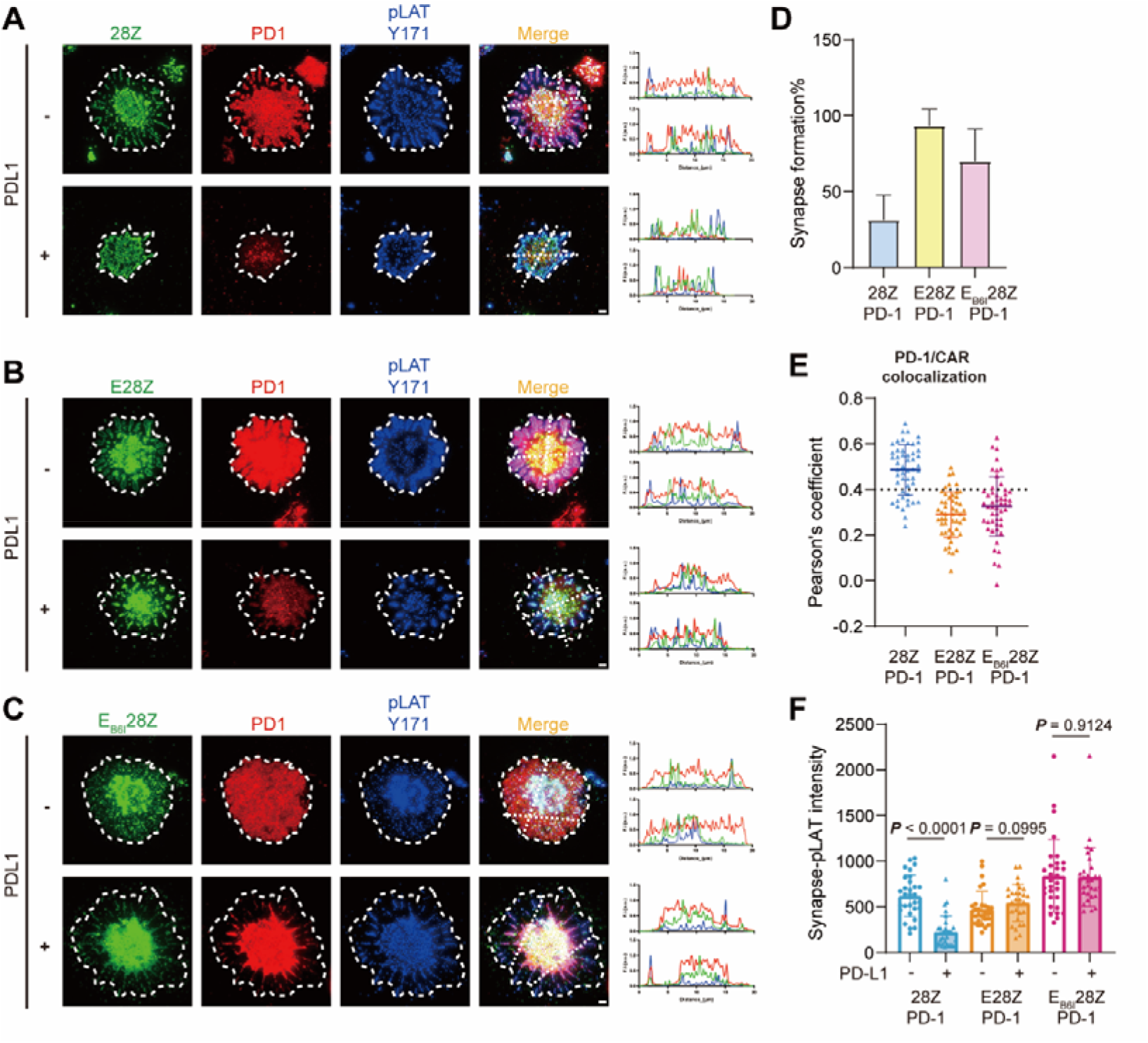
E-CAR counteracts PD-1-mediated immunological synapse and signaling inhibition. **(A-C)** Immunological synapse imaging of 28Z and E-CARs under PD-L1 ligation. Jurkat cells expressing EGFP-tagged anti-CD19 28Z (A), E28Z (B), or E_B6I_28Z (C) CARs together with PD-1-mScarlet3 were stimulated on supported lipid bilayers (SLBs) coated with CD19 (10 nM) in the presence or absence of PD-L1 (10 nM) at 37 °C for 1 h, followed by fixation and intracellular staining for phosphorylated LAT (Y171) with Alexa Fluor 647-conjugated antibody. Representative images of immunological synapses formed by the indicated CAR constructs. Scale bar, 1 μm. (D) Percentage of mature immunological synapses formed by 28Z, E28Z, and E_B6I_28Z CARs. Mature synapses were defined by centralized CAR accumulation, whereas immature synapses displayed dispersed CAR localization at the peripheral ring (n = 10). (E) Colocalization of PD-1 and CAR molecules within the immunological synapse. Colocalization was quantified using Pearson’s correlation coefficient calculated with Fiji (n = 30). (F) CAR downstream signaling, assessed by LAT phosphorylation at Y171. Phosphorylated LAT was quantified as the SD^2^/mean fluorescence intensity within the center of the immunological synapse (n = 30).

### E-CAR T cells preserve potent antitumor functions in the presence of PD-1 signaling

We next examined whether CD3ε integration confers resistance to PD-1-mediated inhibition in primary CAR-T cells. To recapitulate the immunosuppressive microenvironment of solid tumors, we generated anti-MSLN 28Z CAR-T and E-CAR-T cells targeting mesothelin (MSLN), a tumor-associated antigen broadly expressed in multiple solid cancers. MSLN HCT116 colorectal cancer cells, which endogenously express high levels of PD-L1 (PD-L1^+^), together with their isogenic PD-L1-knockout counterparts (PD-L1 null), were used as target cells (**Extended Data Fig. 4A**). Consistent with our biochemical and imaging analyses, PD-L1 expression markedly impaired the cytotoxic activity of 28Z CAR-T cells, whereas E-CAR-T cells largely maintained their antitumor function (**Fig. 6A**). A similar result was observed in an independent CD19 CAR-T model using Raji cells engineered to overexpress PD-L1, which normally express minimal endogenous PD-L1(**Extended Data Fig. 4A and 4B**). We next assessed surface CD69 expression as a marker of T-cell activation. Following co-culture with either wild-type or PD-L1-knockout HCT116 cells, CD69 expression remained comparable in E-CAR-T cells but was markedly reduced in 28Z CAR-T cells upon PD-L1 engagement, indicating that E-CAR preserved activation signaling despite PD-1 ligation (**Fig. 6B**).

**Figure 6.**
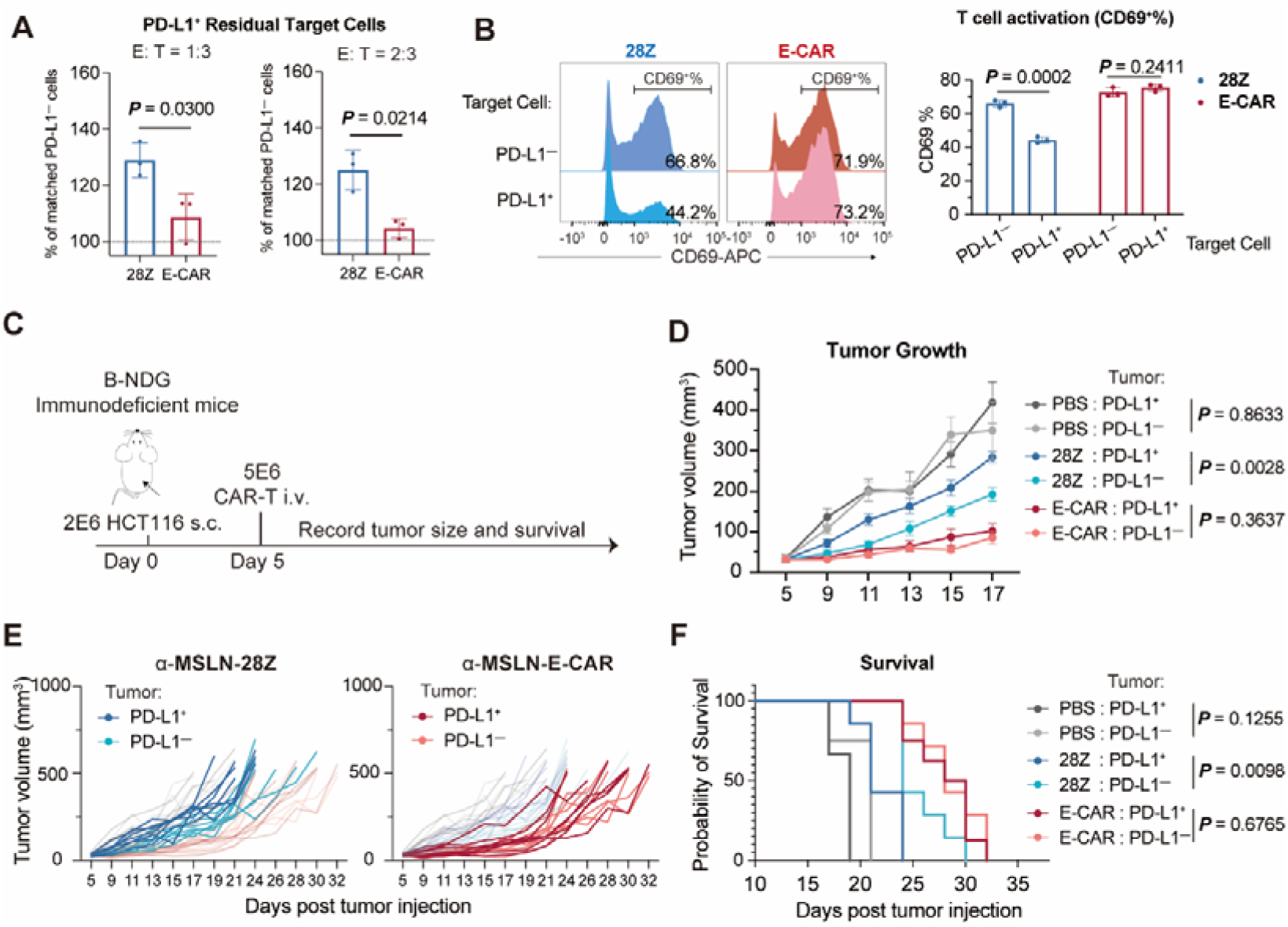
E-CAR-T cells maintain potent antitumor activity against PD-L1-high solid tumors. **(A)** PD-L1-mediated suppression of CAR-T cell cytotoxicity. Normalized PD-L1 HCT116 tumor cell numbers relative to matched PD-L1 target cells following co-culture with anti-mesothelin (MSLN) 28Z or E-CAR (E_B6I_28Z) T cells at the indicated effector-to-target (E: T) ratios (n = 3). CAR-T cells were pre-stimulated for 24 h to induce endogenous PD-1 expression before co-culture with tumor cells. **(B)** Flow cytometric quantification of surface CD69 expression on anti-MSLN 28Z CAR-T and E-CAR (E 28Z) T cells following 4 h co-culture with PD-L1 or PD-L1 HCT116 cells at an E:T ratio of 1:1 (n = 3). PD-1 expression was pre-induced as described in panel (a). **(C-F)** Mesothelin-positive solid tumor model. Immunodeficient B-NDG mice were subcutaneously inoculated with WT (PD-L1 ) or PD-L1 knockout (PD-L1 ) HCT116 cells. Five days after tumor engraftment, mice were administered PBS, anti-MSLN 28Z CAR-T cells, or E-CAR-T cells intravenously. (C) Schematic workflow of the subcutaneous HCT116 adoptive CAR-T cell transfer model. Tumor growth and animal survival were monitored. (D) Mean tumor growth curves for each treatment group, plotted until the first mouse reached the predefined humane endpoint (tumor volume ≥ 500 mm³). Tumor volume was calculated as (width² × length)/2. (E) Individual tumor growth trajectories of mice bearing WT or PD-L1 knockout HCT116 tumors across all treatment groups. (F) Kaplan-Meier survival curves comparing overall survival among treatment groups. Group sizes were: n = 7 for 28Z/PD-L1 , n = 7 for 28Z/PD-L1 , n = 7 for E-CAR/PD-L1 , n = 8 for E-CAR/PD-L1 , n = 4 for PBS/PD-L1 , and n = 3 for PBS/PD-L1 . Data are presented as mean ± standard deviation (SD) for panels (A) and (B), and as mean ± standard error of the mean (SEM) for panel (E). Statistical analyses were performed using two-tailed unpaired t-tests for panels (A) and (B), two-way ANOVA for panel (D), and the log-rank (Mantel-Cox) test for panel (F).

To profile global transcriptional perturbations induced by PD-1 signaling, we performed bulk RNA sequencing of CAR-T cells post-tumor co-culture. Strikingly, only 18 differentially expressed genes (DEGs) were identified between PD-L1-WT and PD-L1-KO conditions in E-CAR-T cells, compared with 396 DEGs detected in 28Z CAR-T cells (**Extended Data Fig. 4D**). KEGG pathway analysis revealed that 28Z CAR-T cells exhibited enrichment of 10 pathways when comparing coculture with PD-L1-knockout versus wild-type HCT116 cells (**Extended Data Fig. 4E**), including pathways associated with T-cell activation (**Extended Data Fig. 4F**). These enriched pathways were supported by multiple DEGs identified in the 28Z CAR-T cells **(Extended Data Fig. 4G)**. In contrast, E-CAR-T cells showed no significant enrichment pathways under the same conditions, with negligible corresponding DEGs abundance (**Extended Data Fig. 4F and 4G**), demonstrating that PD-1 signaling barely remodels E-CAR-T transcriptomic programs.

To evaluate E-CAR T cells-mediated resistance to PD-1 signaling *in vivo*, a solid tumor model using HCT116 cells was established. Immunodeficient B-NDG mice were subcutaneously inoculated with either wild-type (PD-L1^+^) or PD-L1-ablated (PD-L1 null) HCT116 cells, which exhibited comparable growth rates *in vitro*. Five days post tumor implantation, mice were treated with anti-mesothelin 28Z or E-CAR-T cells (**Fig. 6C**). In the 28Z CAR-T treatment group, PD-L1 expression significantly accelerated tumor progression and shortened mouse survival, confirming *in vivo* PD-L1-mediated CAR-T immunosuppression **(Fig. 6D-6F)**. In contrast, E-CAR-T cells exerted robust tumor growth control, and showed marginal tumor progression in response to PD-L1 expression, without compromising long-term host survival (**Fig. 6D-6F**). These findings corroborate that E-CAR-T cells effectively resist TME-intrinsic PD-1 checkpoint suppression and maintain durable solid-tumor antitumor efficacy *in vivo*, highlighting their potential to overcome checkpoint-mediated immunosuppression in solid tumors.

## Discussion

In this study, we established a biomolecular condensation system to investigate signalosome assembly during CAR activation and uncovered an additional layer of CAR signaling regulation. We found that phosphorylated CD28-based CAR (p28Z), but not phosphorylated 4-1BB-based CAR (pBBZ), undergoes LCK-dependent biomolecular condensation, which further recruits unphosphorylated CAR molecules to amplify signalosome assembly. Unexpectedly, we found that the cytoplasmic tail of PD-1 alone was sufficient to compete for LCK binding and disrupt p28Z-LCK condensates independently of SHP1/2, thereby impairing immunological synapse formation. Incorporation of the CD3ε intracellular domain enhanced LCK recruitment, preserved condensate assembly despite PD-1 engagement, and rendered E-CAR resistant to PD-L1-mediated suppression in solid tumor models. Together, these findings identify condensate assembly as an important regulatory layer of CAR signaling and provide a strategy to enhance CAR function in immunosuppressive tumor microenvironments.

Biomolecular condensation has recently emerged as a fundamental mechanism for organizing immune receptor signaling^42^. Phase separation has been implicated in the assembly of TCR, LAT and downstream signaling complexes, enabling rapid signal amplification through local enrichment of signaling molecules^37,42,43^. Our findings extend this concept to engineered antigen receptors by demonstrating that CAR signaling is likewise organized through dynamic condensate assembly. Importantly, we show that LCK and activated CARs not merely form condensates but actively links unphosphorylated CARs, thereby promoting progressive signalosome assembly. The potent signaling capacity of CD28-based CARs may be advantageous in solid tumors^26^, where rapid tumor-cell elimination is often required.

However, solid tumors are characterized by highly immunosuppressive microenvironments enriched in inhibitory checkpoint pathways which in nature inhibit T cell potency^44^. Among these, PD-1 represents one of the major barriers limiting CAR-T efficacy. PD-1 is generally considered to suppress T cell activation through phosphorylation-dependent recruitment of SHP2, leading to dephosphorylation of proximal signaling components^45–47^. Our findings reveal an additional mechanism that precedes phosphatase recruitment, which provides a mechanistic explanation for previous observations that PD-1 remains partially inhibitory even in SHP1/SHP2 double-knockout T cells^48^. Remarkably, the PD-1 cytoplasmic tail alone was sufficient to disrupt CAR-LCK condensates, indicating that checkpoint-mediated inhibition can occur through direct competition for kinase recruitment. Because both CAR activation and PD-1 phosphorylation depend on Src-family kinases^46^, competition for the LCK binding to CAR provides a plausible mechanism by which PD-1 rapidly suppresses signalosome assembly before phosphatase-dependent signal attenuation occurs. These observations expand the current understanding of PD-1 function from a phosphatase-recruiting receptor to a regulator of condensate organization, suggesting that checkpoint receptors may suppress immune signaling by destabilizing higher-order signaling assemblies.

Our findings further demonstrate that enhancing productive LCK recruitment through additional intracellular modules represents an effective strategy to overcome this inhibitory mechanism. Rather than preventing PD-1 engagement, incorporation of the CD3ε intracellular domain shifted the competitive balance toward CAR-associated LCK recruitment, thereby preserving signaling condensate assembly and maintaining downstream signaling under PD-L1-rich conditions. CD3ε has been reported to play a central role in orchestrating LCK recruitment during physiological T-cell activation^49–51^. Multiple intracellular motifs within the CD3ε cytoplasmic domain have been implicated in this process, including the basic residue-rich sequence (BRS), which regulates membrane association and LCK accessibility^52,53^, the RK motif, which directly contributes to LCK recruitment^53,54^, and the proline-rich sequence (PRS)^37,55^, which has also been implicated in promoting productive LCK engagement. Together, these motifs facilitate efficient LCK recruitment to the TCR complex and initiate proximal T-cell signaling. Notably, the E_B6I_28Z construct lacks the PRS motif. Consistent with the reported role of PRS in LCK recruitment, E_B6I_28Z exhibited a modest reduction in resistance to PD-1 incorporation into CAR signaling condensates compared with the full-length CD3ε-containing construct. Nevertheless, E_B6I_28Z remained largely resistant to PD-1-mediated suppression, and proximal signaling, as reflected by preserved LAT phosphorylation following PD-L1 ligation, was sufficiently maintained. However, CD3ε also interacted with negative regulators for T cells, such as LAG3 and Csk. LAG3 has recently been reported to undergo phase separation with the CD3ε-containing TCR/CD3 complex^40,56^, but incorporation of CD3ε into E-CAR did not promote LAG3 incorporation or disrupt LCK-pE-CAR/ condensates. One possible explanation is that, the CD28 and CD3ζ signaling motifs within E-CAR provide additional binding interfaces for LCK, creating a condensate environment that favors productive LCK engagement. Similar to the competition observed in the PD-1 context, these additional LCK-CD28 interactions may offset LAG3 binding to CD3ε, thereby preserving condensate integrity despite the presence of inhibitory receptors. Similarly, Csk was also unable to disrupt pE-CAR/LCK condensates, likely owing to the additional LCK recruitment mediated by the CD28 and CD3ζ signaling domains. These observations support a design principle that we term modular counterplay, whereby intracellular signaling modules do not function independently but instead dynamically compete and cooperate for shared signaling components within biomolecular condensates. Rather than simply adding stimulatory signaling capacity, appropriately selected modules can simultaneously reinforce productive molecular interactions while counterbalancing inhibitory ones, thereby generating synergistic signaling outputs that exceed the simple sum of the individual signaling modules. More broadly, engineering modular counterplay may represent a general strategy for designing next-generation CARs with enhanced resilience to checkpoint-mediated suppression and improved efficacy against solid tumors.

Within the tumor microenvironment (TME), PD-L1 expressed by both tumor cells and myeloid cells constitutes a major barrier to CAR-T cell function by driving PD-1-mediated suppression^39,57^. Accordingly, blockade of the PD-1/PD-L1 axis has demonstrated durable clinical benefit in patients with PD-L1-positive tumors, and sequential combination of CAR-T therapy with immune checkpoint blockade has been shown to reduce disease recurrence, delay progression, and overcome therapeutic resistance in high-risk settings^58–60^. Likewise, armored CAR-T cells engineered to locally secrete PD-1 antibodies or nanobodies have exhibited improved antitumor efficacy, further highlighting the importance of relieving PD-1-mediated inhibition^56,61^. Together, these findings underscore that targeting PD-1 signaling is an effective strategy to enhance CAR-T cell function. Nevertheless, systemic PD-1 blockade remains dependent on optimized dosing and treatment schedules, and may increase the risk of excessive immune activation, including cytokine release syndrome (CRS)^59^, as well as immune-related adverse events and hyperprogression in certain contexts^62–65^. In contrast, E-CAR represents a streamlined and intrinsically self-regulated engineering solution that counteracts PD-1-mediated suppression by rewiring CAR signaling at the receptor level, thereby minimizing reliance on exogenous checkpoint blockade and simplifying therapeutic implementation, particularly in the context of solid tumors.

Collectively, our study identifies a previously unrecognized mechanism by which the PD-1 cytoplasmic tail suppresses CAR signaling through disruption of LCK-driven biomolecular condensates, independently of its canonical phosphatase-dependent pathway. More importantly, we establish modular counterplay as a conceptual framework for CAR engineering, whereby intracellular signaling modules dynamically compete and cooperate for shared signaling components to reshape condensate composition and signaling output. Our findings suggest that exploiting modular counterplay will provide a general strategy for engineering next-generation CARs and other synthetic immune receptors with enhanced resilience to checkpoint-mediated suppression and improved therapeutic efficacy in solid tumors and other immunosuppressive diseases.

## Acknowledgements

We thank Qing Bian and Yun Feng for the help in TIRF-SIM imaging and data processing with Imaris, thank Junying Jia and Shu Meng (Core Facility, Institute of Biophysics, CAS) for technical support in the flowcytometry analysis, and thank Xiang Ding and Mengmeng Zhang for LC-MS/MS technical support in protein phosphorylation analysis. All the imaging experiments were performed at the Center for Biological Imaging (CBI), Institute of Biophysics, Chinese Academy of Science. This work was supported by grants from the National Natural Science Foundation of China (32090044 to J. L., T2394512 and 32200549 to H. C.), the Strategic Priority Research Program of the Chinese Academy of Sciences (XDB1000103 to J. L.), and Beijing High Innovation Plan (202504841084 to J.L.). C.X. is funded by the Strategic Priority Research Program of the Chinese Academy of Sciences (XDB0990000), and the Shanghai Pilot Program for Basic Research-Chinese Academy of Sciences, Shanghai Branch (JCYJ-SHFY-2022-009). C.X. is also a scholar of the Shanghai Academy of Natural Sciences (SANS).

## Author contributions

J.Z.L. conceived the project and supervised all experiments. C.X. contributed to the project design and co-supervised the cellular and animal experiments. H.C. contributed to the experimental design of CAR condensation. Y.J. performed protein purification, condensation assays, and immunological synapse imaging experiments. Z.R. performed the human primary CAR-T cell experiments and tumor studies. J.J.L. performed protein purification and construct design. H.Y. performed the bioinformatic analyses. X.L. generated the PD-1/PD-L1 plasmids, SHP1/SHP2-double-knockout cells and contributed to discussions. Y.J. wrote the manuscript. Z.R., J.Z.L. and C.X. revised the manuscript.

